# Group size planning of breedings of gene-modified animals

**DOI:** 10.1101/2021.09.17.460764

**Authors:** Vladislava Milchevskaya, Philippe Bugnon, Emiel B.J. ten Buren, Frank Brand, Achim Tresch, Thorsten Buch

## Abstract

Animal breeding is time-consuming, costly, and affected by stochastic events related to Mendelian genetics, fertility, and litter size. Careful planning is mandatory to ensure a successful outcome using the least number of animals, hence adhering to the 3Rs of animal welfare. We have developed an R package, accessible also through an interactive website, that optimizes breeding design and provides a comprehensive report suitable for any breeder of genetically defined traits.

## Main

Rodent research and hence breeding has undergone an explosive development. Mouse Genome Informatics counts 47’000 curated and 14’000 non-curated entries for mouse lines^1^. A total of 7 Mio rodents per year are used for creation and maintenance of gene-modified lines in the European Union alone^2^. In most projects not only single mutants are bred and analyzed: rather combinations of multiple alleles of different genes have become mainstay in research. These complex genotypes are assembled through time and cost intense breeding schemes^1^.

The practice of killing animals because they do not carry specific traits or are not needed has become under scrutiny^3,4^, in laboratory animal science^5^, farming^6,7^, and zoos^8–11^.

The causes of unwanted surplus in laboratory animal facilities were identified, among them: genetics of breeding, sex preference and the inability to match supply with demand^12^. Stochastic fluctuations in allele distribution, in fertility (some breeding pairs will produce no offspring), and litter size (number of pups born/weaned per litter) have a large influence on breeding outcomes. Their neglect results in unnecessary breeding delays and scientifically unjustified animal use. We describe here software that enables researchers to plan their breedings with a given success probability, integrating the stochastic effects caused by Mendelian genetics, fertility and litter size. Typically, genotype frequencies in offspring are obtained via Punnett square^13^ (Fig 1A). However, in a single or few litter(s), the observed frequencies may differ substantially due to random fluctuations. Given a fixed litter size, the number of e.g. -/- animals being present in a litter from +/-parents follows a Binomial distribution (Fig 1B, Mendelian probability c=0.25). If the breeding outcome does not follow classical Mendelian frequencies (e.g. embryonal deaths^14,15^), the probabilities of occurrence change (Fig. 1B, c = 0.2). Furthermore, litter size itself is a variable that can either be positive (number of offspring in a litter when the mating is successful) or turn zero when the mating is unsuccessful. We collected the empirical distributions of litter sizes for 8 mouse lab strains and found that most of them could be approximated well by a Poisson distribution (Figure 1C, Suppl. Figure 1), when unsuccessful matings are excluded. The fraction of successful matings (fertility) in our calculator is obtained from the values reported by the Jackson lab (http://www.informatics.jax.org/silver/tables/table4-1.shtml). Having specified these components, we derived the distribution of the target offspring number as a function of the number of matings (Figure 1D, Supplementary Methods). The probability of successfully obtaining the desired number of pups of needed genotypes from a specific setup can then be quantified. In the 1980s, Festing proposed a method for modelling the probabilistic outcomes in fertility and litter size^16^. We developed an alternative method that can be proven more accurate (Supplementary Methods) and thereby often reduces the number of matings to be set up (Fig 2A). We also show that the simplistic use of the expected target animal number derived from Mendel’s laws combined with average litter size underestimates the required number of matings dramatically (Fig. 2A, Mendel). We recommend setting the desired success probability not overly high (e.g. below 0.95) since further confidence increase is costly in terms of additional matings. (Figure 2B, Suppl. Figure 3).

**Fig. 1.**
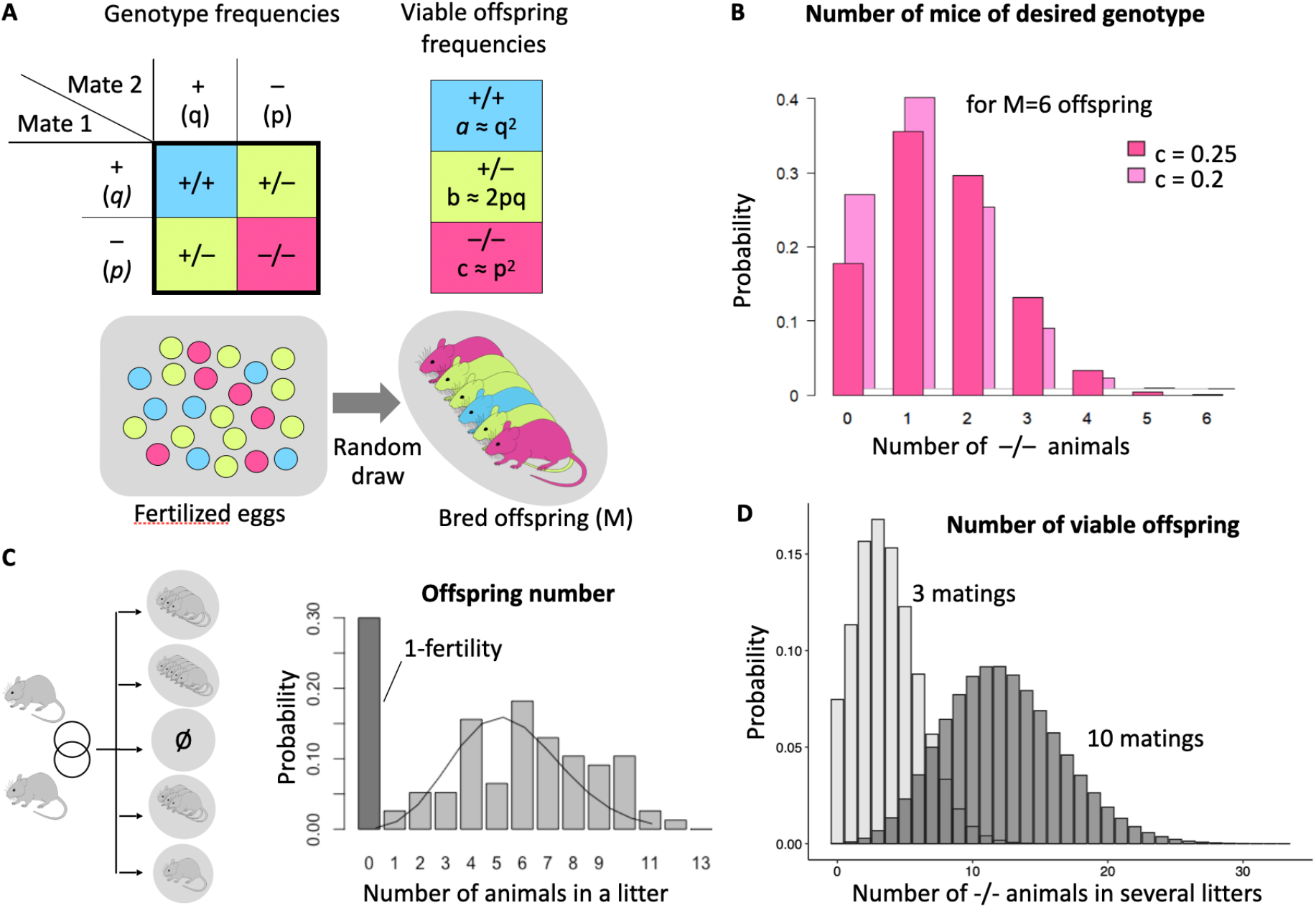
Stochastic Effects in Breeding. (A) Mendelian model of the outcome of a single breeding of two heterozygous animals. + and - denote different gene alleles occurring with probabilities *q* and *p*, correspondingly. Theoretical frequencies of the +/+, +/- and -/- offspring are q^*2*^, *2pq* and p^*2*^. An outcome of single breeding is depicted as a random draw from the theoretical distribution above. (B) Probability that out of 6 offspring exactly X animals will have the desired -/- genotype, given the probability *p*^*2*^ of the -/- genotype is 0.25 or 0.2. (C) Example of the number of animals born out of a single breeding when fertility is taken into account: There is a non-zero chance that 0 animals are born (1-fertility). (D) Distribution of -/- animals born for a genotype frequency of 0.25, for 10 (grey) respectively 3 (light grey) breedings.

**Figure 2.**
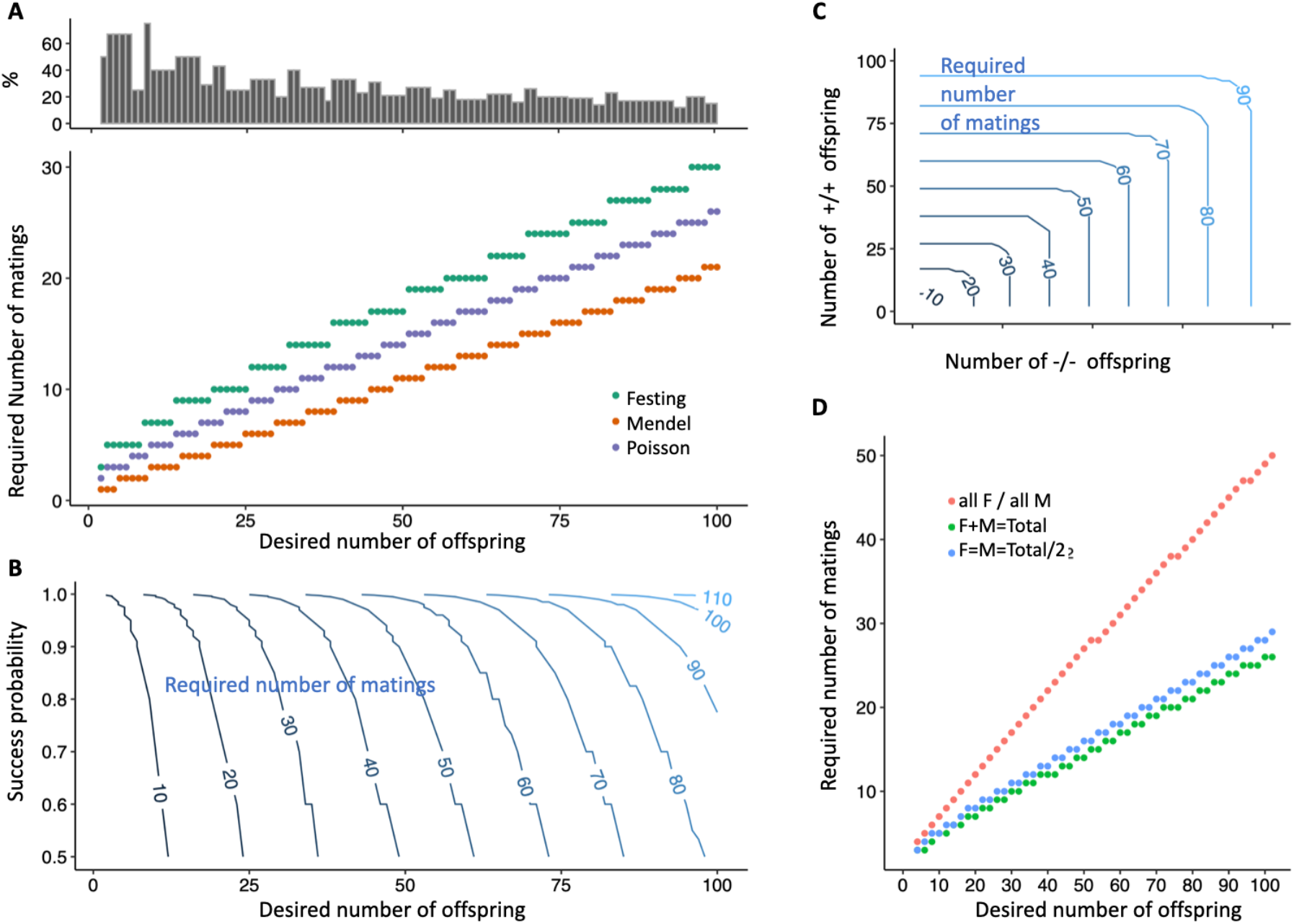
Performance of breeding models, in the case of an average litter size of 7 and a mouse fertility of 70%. (A) Bottom: Minimal required number of breedings (Y-axis) needed to obtain the desired number of offspring (X-axis) with 90% confidence, as calculated by three methods: The naïve expectation due to Mendelian frequencies (see Suppl.), the gold standard textbook model suggested by Festing, and our method, denoted as Poisson. Top: Relative surplus of breedings (in %) required by the textbook method (Festing), measured against our method. (A, bottom) (B) Minimal number of breedings needed to obtain a certain number of offspring (X-axis) with a defined probability of success (Y-axis). (C) Minimal number of breedings (contour lines) required in a setup where groups with two different genotypes need to be produced by the same breedings. (D) Minimal number of breedings required to obtain offspring of specific sexes with 90% confidence, given that both female (F) and male (M) pups are born with equal probability. Shown are three scenarios: X offspring required to be of the same sex required (red), X offspring of any sex required (green), X offspring with balanced cohorts of each sex required, i.e. X/2 male and X/2 female pups (blue).

Often multiple genotypes need to be produced by the same set of breeding pairs, e.g., identical numbers of +/+ and -/- animals from +/- parents. Such a setup reduces success probability further and requires additional matings (Figure 2C). The same calculations apply to group size planning for obtaining defined numbers of animals of either sex. While some experimental designs require all animals to be of the same sex, alternative designs can include both sexes (and account for sex-specific effects)^17^. A group size planning for inclusion of both sexes at identical number increases the required matings only slightly over simple use of all males and females born, without fixed ratio (Figure 2D). However, the use of only one sex increases the required number of matings massively^17^ (Figure 2D).

To facilitate appropriately powered breeding for the practitioner, we incorporated the algorithms and data for sample size calculation into the R package ’*BreedingCalculator*’, available at the GitHub repository https://github.com/VladaMilch/breedingCalculator. Simplified, interactive access to this package is provided on the website https://www.ltk.uzh.ch/en/Breeding.html. We complemented the package with an additional parameter, namely effective fertility, based on the numbers given by Festing^16^. Effective fertility comes into effect when the age of the experimental cohort is fixed to a short time period such as birth within 1, 2 or 3 days^16^.

Optimization of breeding protocols for reduction of animal use is an ethical obligation mandated within the commonly applied 3R (replace, reduce, refine) principle. Yet, the very basic biology of mammalian genetics and associated stochastic processes inevitably create surplus animals without further use in either experiments or breeding. We have developed an R package that supplies the optimal solution, i.e. the least number of required animals, for every use case described above. It uniformly improves over previously published tables and schemes (e.g., https://www.jax.org/jax-mice-and-services/customer-support/manuals-posters-and-guides/jmcrs-manuals-guides). We removed from the workflow any form of guess work commonly done by scientists to adjust for self-experienced stochastic effects, colloquially referred to as *Murphy’s law*. Also, through appropriate group size calculations for breedings, experiments are more likely to be conducted as planned, which serves their reproducibility. It may seem that powered breeding planning increases the number of animals produced for an experiment. But this is not the case, because if the planning is not adequate and the desired number of animals is not reached, a new breeding round will be required and the animals from the first round remain unused. Hence, correct planning allows in a precise way to reach the required number of animals by reducing the number of unsuccessful breeding attempts. While unequal use of the sexes in animal experimentation has been discussed thoroughly, with statistical solutions regarding experimental designs suggested, we here provide evidence that restricting experiments to one sex unnecessarily leads to additional breedings and hence unused offspring beyond a simple doubling. Taken together, we provide a method supported by software to minimize animal numbers in breeding while explicitly controlling for experiment’s success, being in full compliance with the 3R rule.

## Supplementary Figures

**Figure S1:**
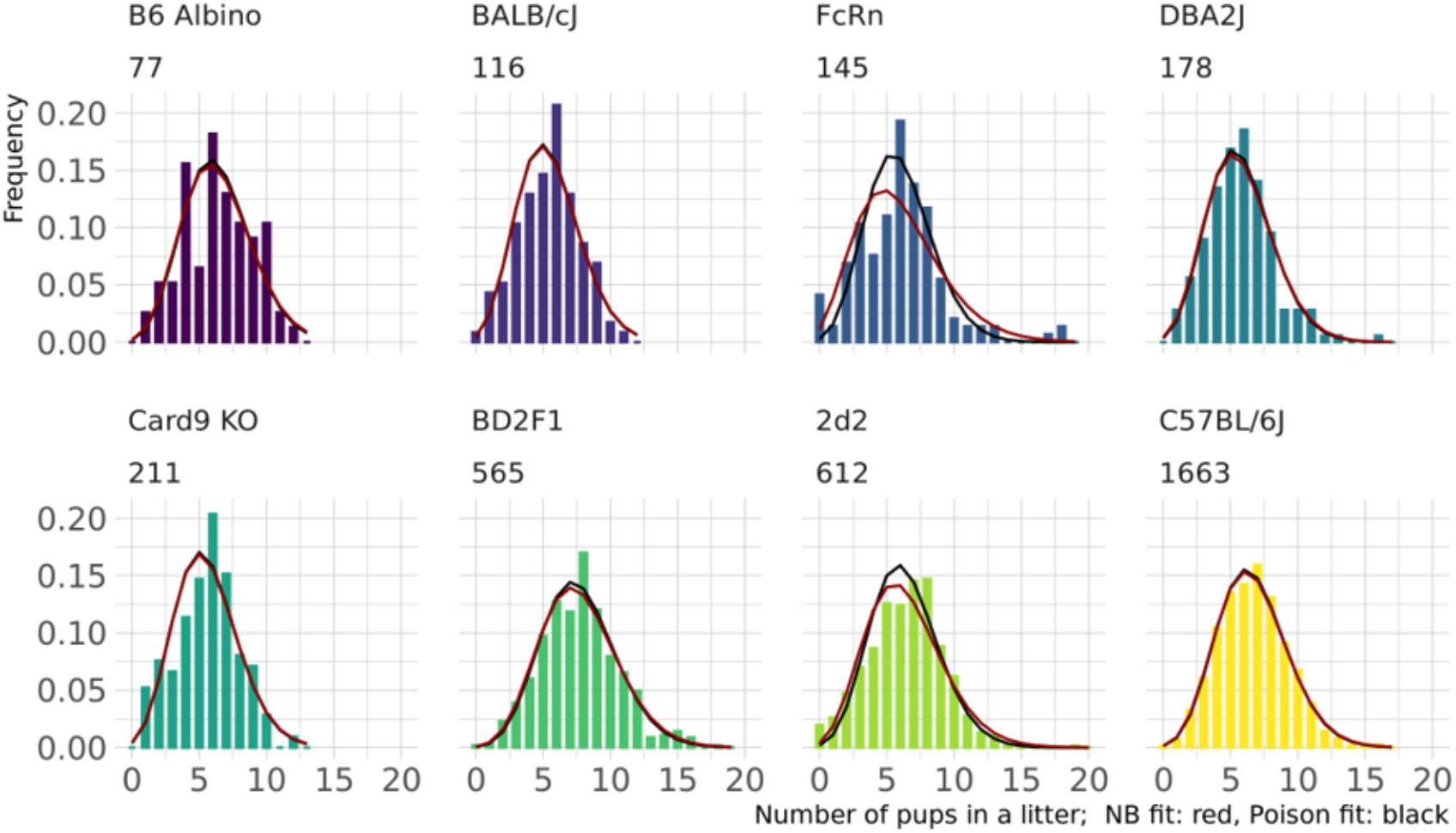
Litter size distribution for different mouse strains, estimated from the corresponding number of litters (under the strain name). On top of each histogram, fits of the Negative Binomial (red) and Poisson (black) distributions are displayed. The fits were performed using the Maximum Likelihood approach implemented in the MASS package in R. Strain/line names are indicated and refer to: B6 Albino - B6N-Tyrc-Brd/BrdCrCrl, BD2F1 - B6D2F1, Card9 KO – C57BL/6N-Card9^em1ltk^, FcRn – C57BL/6.Cg-Fcgrt^tm1Dcr^ Tg(FCGRT)32Dcr/DcrJ, 2d2 -Tg(Tcra2D2,Tcrb2D2)1Kuch, total number of used litters is indicated under the line name.

**Figure S2:**
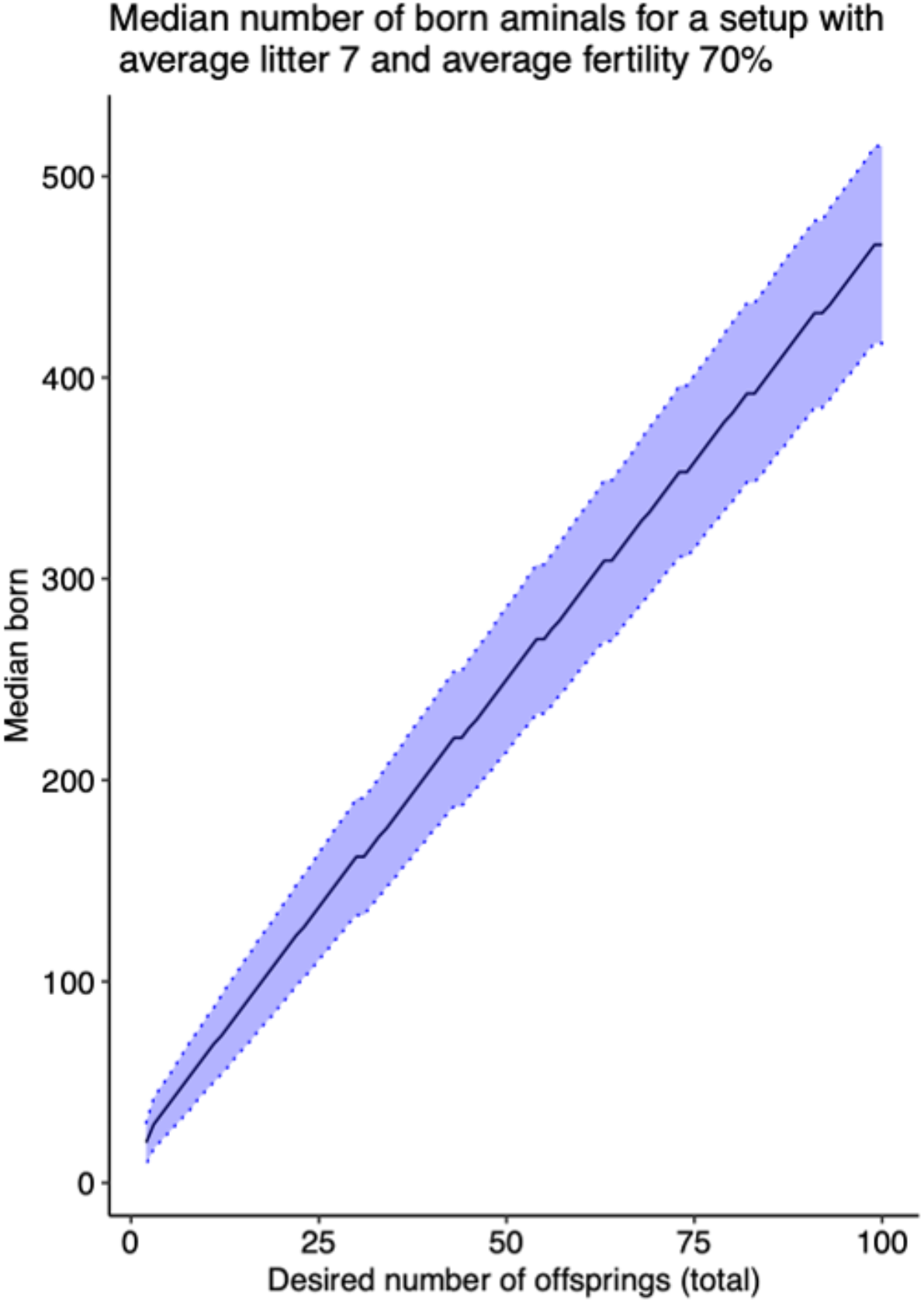
Total number of born animals, when the breeding setup is designed to yield no less than X animals with success probability 90%. Blackline depicts the median number of animals born, the blue area around corresponds to the symmetric confidence interval of 0.8.

**Figure S3:**
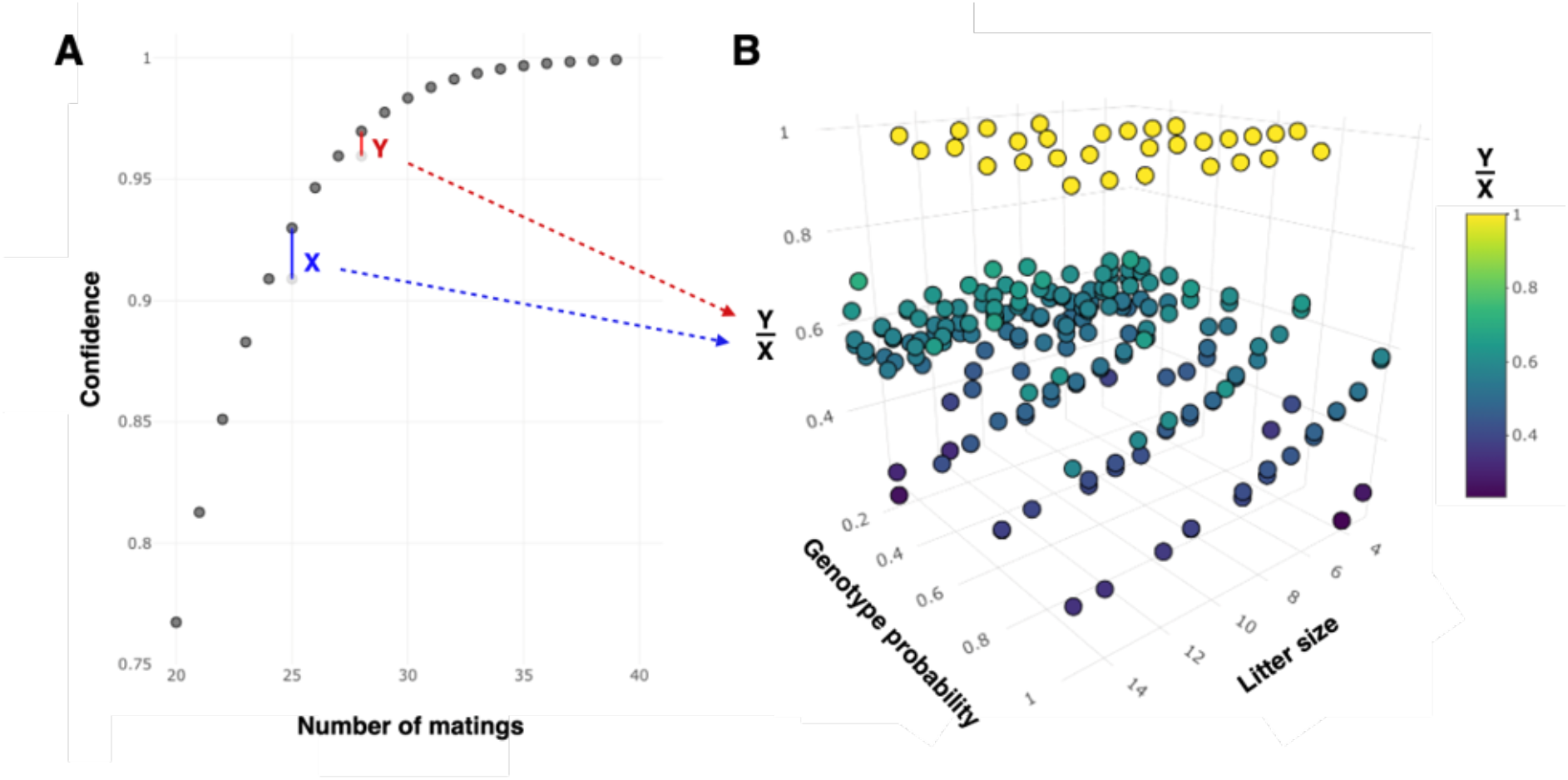
Optimal confidence level choice. Panel A shows an example of the relation between the number of matings and the confidence in the successful outcome of the breeding; segments X (blue) and Y (red) show the confidence increase granted by one additional mating: X – starting from the number of matings sufficient for the 90% confidence, Y – from the number of matings sufficient for the 95% confidence. One can see that the higher the target confidence, the smaller the confidence increase – which is especially pronounced for the high confidence values (flat curve). The parameters used for this example: the average litter size of 5, fertility of 50%, desired genotype probability of 50%, the desired number of pups of this genotype – 20. Panel B shows the ratio of the 95% and the 90% confidence increases from panel A, calculated for different starting parameters for the litter size, the fertility (not shown) and the desired genotype probability. Hence, one point corresponds to a single example as in panel A. For some cases (yellow), when very few matings are required to yield a successful breeding, there is no difference between X and Y, thus the ratio is 1. However, for the majority of breeding parameters, the increase in confidence above from 95% is from 1.5 to 10 fold smaller than the increase from 90% (Y/X values below 0.6). We therefore recommend choosing the target confidence level not too high (often, below 95%).

## Notes

### Competing Interest Statement

The authors have declared no competing interest.

