## Supplementary Methods for "Group size planning of breedings of gene-modified animals"

August 2021

#### 1 Notations and Problem Statement

##### 1.1 Total number of offspring

Let us assume that a researcher conducts $M$ matings, i.e. introduces $M$ female animals each to their respective male for breeding. Further, we use the following notations.

Let $\xi_i, \;\; i \in \{1, \ldots, M\}$ be a non-negative integer random variable representing the number of offspring born of the $i$-th mating and

$$P(\xi_i = x) = \begin{cases} 1 - p_f & : x = 0 \\ \textit{some distribution} & : x > 0 \end{cases} \tag{1}$$

where $p_f$ is the fertility – the probability that a mating was successful, namely $p_f = P(\xi_i > 0)$, – which is a property of the animal strain, i.e. it is the same for all matings $i \in \{1, \ldots, M\}$.

Further, let $N_d \in \mathbb{N}$ be the desired number offspring to be born, and let a random variable $\chi = \sum\limits_{i=1}^{M} \xi_i$ denote the actual number of offspring born. Then, the confidence that the breeding experiment yields at least $N$ offspring (success confidence) is as follows: $P_d = P(\chi \geq N)$.

If the distribution of $\xi_i$ is given, the distribution and the corresponding confidence intervals of $\chi$ can be derived as a convolution of $M$ i.i.d. variables – either numerically or analytically. Therefore, the above confidence $P_d$ can be represented as a function of the number of matings, i.e.

$$P_d = P_d(M) = P\Big(\sum_{i=1}^{M} \xi_i \geq N\Big). \tag{2}$$

Now we can formulate the goal of our breeding set-up in the above notations.

**Problem 1** *For a desired confidence level $p_{success} < 1$, number of offspring $N \in \mathbb{N}$ and a given distribution of the i.i.d. $\xi_i \;\; i = 1, 2, ...,$ determine the smallest $M$ that guarantees $P_d(M) \geq p_{success}$, i.e. find the*

$$\hat{M} = \arg\min_{M \in \mathbb{N}} P\Big(\sum_{i=1}^{M} \xi_i \geq N\Big) \geq p_{success}. \tag{3}$$

Alternatively, we can re-write eq. (5) in this form (will be used later):

$$\hat{M} = \arg\min_{M \in \mathbb{N}} P\Big(\sum_{i=1}^{M} \xi_i < N\Big) < 1 - p_{success}. \tag{4}$$

##### 1.2 Offspring of multiple genotypes

The previous problem covers those breeding set-ups where the *total* offspring number is of interest. Another common requirement is for offspring of multiple genotypes to be born from the same breeding set-up.

Let us consider a breeding, in which offspring of $G > 1$ different genotypes can be born. The probability of a single offspring to belong to one of the $G$ genotypes is known and denoted as $\{p_1, ..., p_G\}$, $\sum_{g=1}^{G} p_g = 1$. Further, for each genotype $g \in \{1, ..., G\}$, the researcher desires to get a certain number $n_g \geq 0$ of offspring.

Similarly to the previous notation, we denote the actual number of genotype $g$ offspring born from a single mating as $\{\xi_i^g\}$, and their sum over all matings as $\chi_g = \sum_{i=1}^{M} \xi_i^g$. For a breeding to be successful, we need to ensure that $\chi_g \geq n_g$ for each $g \in \{1, ..., G\}$.

**Problem 2** *Here we assume that the total numbers of offspring from each mating $\xi_i$ $i = 1, 2, ...$ are i.i.d and come from a known distribution, and the genotype probabilities are $\{p_1, ..., p_G\}$. For the desired numbers of offspring of each genotype $(n_1, ..., n_G)$ and a desired confidence level $p_{success} < 1$, determine the smallest $M$ that guarantees the breeding to be successful with probability no less than $p_{success}$, i.e.*

$$\hat{M} = \arg\min_{M \in \mathbb{N}} P\Big(\chi_g \geq n_g \text{ for all } g \in \{1, ..., G\}\Big) \geq p_{success}. \tag{5}$$

#### 2 Festing sample size calculation

In the textbook method proposed by Festing [1], Problem 1 is addressed using certain assumptions:

1. $p_{success} = 0.95$
2. the mean litter size $\mu$ and the effective fertility $p_f$ of an animal strain are given
3. $P(\xi_i = x | x > 0) \sim N(\mu, \delta = 2.5)$

The last assumption may seem odd now, since an integer value (the number of born offspring) is modeled with a continuous distribution. However, in the 1980-th it was not possible to calculate convolution of any distribution numerically. Festing et al. proposed a remedy for it: due to the central limit theorem, the sum of sufficiently many random variables will have an approximately Gaussian distribution. Moreover, researchers could look up quantiles of a Gaussian distribution in a specialized directory. Therefore, assumption (3) makes a reasonable prediction when the total number of offspring born from *many* matings is concerned.

Further, let $l = l(M)$ be the number of non-empty litters produced by $M$ matings, i.e. the number of such $i$ that $x_i > 0$. Consider a situation when $M$ mating produced exactly $L$ litters have. Given assumption (2), $l(M)$ will follow a Binomial distribution, i.e.

$$P(l(M) = L) = Bin(L; M, p_f).$$

Therefore, if we denote the Gaussian cumulative probability function as $P_N(\cdot)$ and the Binomial cumulative probability function as $P_{Bin}(\cdot)$, eq. (4) can be viewed as follows:

$$\begin{aligned}
P\Big(\sum_{i=1}^{M} \xi_i < N\Big) &= \sum_{L=0}^{M} P(l(M) = L)\ P\Big(\sum_{l=1}^{L} (\xi_i | \xi_i > 0)\ < N \mid l = L\Big) \\
&= \sum_{L=0}^{M} P_{Bin}(l = L; M, p_f)\ P\Big(\sum_{l=1}^{L} (\xi_i | \xi_i > 0)\ < N \mid l = L\Big) \\
&\overset{\text{(a)}}{\approx} \sum_{L=0}^{M} P_{Bin}(l = L; M, p_f)\ P_N(\chi < N;\ \mu L,\ 2.5\sqrt{L})\ = \qquad (6) \\
&= \sum_{L=0}^{L_1} P_{Bin}(l = L; M, p_f)\ P_N(\chi < N;\ \mu L,\ 2.5\sqrt{L})\ + \\
&\quad \sum_{L=L_1}^{M} P_{Bin}(l = L; M, p_f)\ P_N(\chi < N;\ \mu L,\ 2.5\sqrt{L})
\end{aligned}$$

For any value $L_1 \in \{1, \ldots, M-1\}$, the last sum can be split into two as below.

$$
\begin{aligned}
&\sum_{L=0}^{M} P_{Bin}(l = L; M, p_f)\ P_N(\chi < N;\ \mu L,\ 2.5\sqrt{L})\ = \\
&= \sum_{L=0}^{L_1} P_{Bin}(l = L; M, p_f)\ P_N(\chi < N;\ \mu L,\ 2.5\sqrt{L}) + \\
&\quad + \sum_{L=L_1}^{M} P_{Bin}(l = L; M, p_f)\ P_N(\chi < N;\ \mu L,\ 2.5\sqrt{L}) \\
&\overset{(a)}{\leq} \sum_{L=0}^{L_1} P_{Bin}(l = L; M, p_f)\ + \\
&\quad + P_N(\chi < N;\ \mu L_1,\ 2.5\sqrt{L_1}) \sum_{L=L_1}^{M} P_{Bin}(l = L; M, p_f)\ \leq \\
&\overset{(b)}{\leq} P_{Bin}(l \leq L_1; M, p_f) + P_N(\chi < N;\ \mu L_1,\ 2.5\sqrt{L_1})
\end{aligned}
\tag{7}
$$

The last expression allows Festing to use the following, computationally very simple, two-step procedure:

1. How many litters $L$ would one need to ensure that the total number of offspring (distributed as $N(\mu L, 2.5\sqrt{L})$) will be larger or equal to $N$ with probability no less than $1-(1-p_{success})/2 = 0.975$? Namely, we determine the smallest suitable $\tilde{L}$ using the quantile of the Gaussian distribution:

$$\tilde{L} = \arg \min_{L \in \{1,\dots,M\}} \quad P\Big(N(\mu L, 2.5\sqrt{L}) \geq N\Big) \geq 0.975$$

2. Further, how many matings would one need to ensure $\tilde{L}$ or more litters (distributed as $Bin(L; M, p_f)$) with probability no less than $1-(1-p_{success})/2 = 0.975$? Namely,

$$\tilde{M} = \arg \min_{M \in \mathbb{N}} \quad P\Big(Bin(M, p_f) \geq \tilde{L}\Big) \geq 0.975$$

Then, we arrive at the following solution to Problem 1:

$$\tilde{M}_{Festing} \approx \arg \min_{M \in \mathbb{N}} P\Big(\sum_{i=1}^{M} \xi_i \geq N\Big) \geq 0.95 \tag{8}$$

#### 2.1 Festing's calculation is approximate

Assumption 3. $P(\xi_i = x | x > 0) \sim N(\mu, \delta = 2.5)$ of the Festing's calculation demands clarification. Though it is unrealistic to assume that the number of offspring in a single litter follows a Gaussian distribution (a continuous distribution with negative values included in the support) due to the LT gives a good approximation for a large sum of litters (say, $\geq$ 10 litters). In this situation, we don't expect a large error to come from the approximate equality (a) in equation (6). Nevertheless, one should still be cautious when only a few litters are summed (can happen when few pups are needed or when the fertility for a mouse strain is low).

Another source of inaccuracy is the inequalities (a) and (b) in equation (7). These upper bounds are rather crude and often leads to $\tilde{M}_{Festing}$ noticeably overestimating the optimal number of matings for Problem 1 (we show this with practical examples in the main text). However, these workarounds were necessary in the 80s, since one could not compute a convolution of several distributions numerically.

Though Festing's estimate may demand more animals than necessary, in most cases it still guarantees the required confidence $p_{success}$ for the number of desired offspring $N$. We thus use it as a gold standard to improve upon.

### 3 Rule based solely on Mendelian expectation for sample sizes

Here we describe what we believe is an intuitive approach to sample the size calculation, to get $N$ offspring in total. [2] Average litter size ($\mu$) and fertility ($p_f$) guide the required number of matings' calculation, i.e.

$$\hat{M}_{mendel} = \frac{N}{\mu \; p_f}. \tag{9}$$

If one requires $n_g$ offspring of a certain genotype $g$ (with its respective probability $p+g$) and no offspring no other genotypes, the number of matings is

$$\hat{M}_{mendel} = \frac{N}{\mu \; p_f \; p_g}. \tag{10}$$

### 4 Our method

First of all, we collected data from different mouse strains to fit the distribution of $\xi_i|\xi_i > 0$. Unfortunately, since unsuccessful matings are rarely recorded, it was practically impossible to estimate the fertility $p_f = P(\xi_i = 0)$ from the data we had. Thus, we used publicly available values of mouse fertility (be strain). For the successful matings, we found that the litter size ($\xi_i|\xi_i > 0$) of most mouse strains can be accurately approximated by a Poisson distribution (see the main text and Fig. S1). The complete distribution of $\xi_i$ is then a zero-inflated Poisson. However, to retain the flexibility, we implemented the calculations for any non-negative integer distribution for $\xi_i$. The only restriction is that the $\xi_i, i = 1, ..., M$ are i.i.d.

In case of multiple genotypes, we assume that a pup is assigned its genotype with probability $\{p_1, ..., p_G\}$, $\sum p_g = 1$, therefore,

$$\xi_i^g | \xi_i \sim Multinom(\xi_i, p_1, ..., p_G), \quad g \in \{1, ..., G\} \tag{11}$$

Then, conditioned that the total number of offspring $\chi = N$, the total numbers of pups per genotype $\bar{\chi}_g = \{\chi_g\}_{g \in \{1,...,M\}}$ also follows a multinomial distribution, namely,

$$\begin{aligned} &\chi = \sum_{i=1}^{M} \xi_i, \quad \chi_g = \sum_{i=1}^{M} \xi_i^g \\ &\bar{\chi}_g|_{\chi=N} \sim Multinom(N, p_1, ..., p_g). \end{aligned} \tag{12}$$

Therefore, for implementation we use the following

$$\begin{aligned} &P(\chi_1 \geq n_1, ..., \chi_G \geq n_g) = \\ &\sum_{N=0}^{\infty} P(\chi = N) P(\chi_1 \geq n_1, ..., \chi_G \geq n_g | \chi = N), \end{aligned} \tag{13}$$

Here $P(\chi_1 \geq n_1, ..., \chi_G \geq n_g | \chi = N)$ is the multinomial distribution, and $P(\chi = N)$ is a convolution of $M$ i.i.d. non-negative integer distributions, which we calculate numerically using the 'distr' R-package. The upper limit for the sum in equation (13) is finite: Either because the empirical distribution for the size of a single litter $\xi_i$ is defined on a finite support, or, in case of Poisson distribution for $\xi_i$, we use a finite support that covers at least $1 - 0.1^{10}$ probability.

#### 4.1 Comparison to the Festing's calculation

Earlier we have considered two approximations made in the Festing's model: the single litter distribution (see (a) in equation (6)), and the work-around to compute a convolution of multiple random variables (see (a) and (b) in equation (7)).

Here, instead of a Gaussian distribution for the litter size, we use a data-proven Poisson fit. This yields accurate results even for experiments with low offspring demand. Further, we calculate the convolution of multiple distribution numerically, which allows us to bypass the use of inequalities (a) and (b) in equation (7), and thus avoid the overestimation.

### References

[1] MFW Festing. Animal production and breeding methods. *UFAW handbook on the care and management of laboratory animals/edited by Trevor B. Poole; editorial assistant, Ruth Robinson*, 1987.

[2] Reginald Crundall Punnett. Mendelism in relation to disease, 1908.
